# Method for determining of cytotoxicity based on the release of fluorescent proteins

**DOI:** 10.1101/2024.08.31.610658

**Authors:** Dmitry Lifanov, Dulamsuren Zorigt, Evgenya Shabalina, Abdullah Khalil, Konstantin Gorbunov, Elena Petersen

## Abstract

This paper describes a method for determining the cytotoxicity of chemical compounds based on the detection of fluorescent proteins - in this case, green fluorescent protein (GFP) and red fluorescent protein (RFP), which are released into the medium from dead cells. This method is similar in principle to the lactate dehydrogenase test (LDH test), but it does not require a reaction with a chromogenic substrate. This method also makes it possible to independently determine the viability of different lines when used in cocultures. Experiments were preformed on a classical monolayer, spheroids and 3D cultures in alginate hydrogel. Capecitabine was used as a model cytotoxic agent. We included liver cells (Huh7) in a coculture model and determined changes in the cytotoxicity levels of capecitabine against NCI-H1299 cells. The experimental part also found that there were differences in sensitivity to capecitabine depending on the type of 3D cultures used.

## Introduction

Currently, many methods are used to determine the cytotoxicity of various chemicals and biological products. They can be classified by performance (screening versus single samples), applicability to cell culture type, and underlying principles. The most widely used screening tests are MTT and LDH, the most widely used methods applicable to individual samples are PI and trypan blue staining. All these methods have certain limitations and advantages.

Unfortunately, there is no single method that allows one to reliably determine TC50/EC50 with good inter-methods convergence. Based on our literature review, many commonly used methods exhibit some bias when compared with other methods[1]. Sometimes these differences are twofold or more[2]. In other cases, this bias may be caused, presumably, by the contribution of the drug itself to the absorption spectrum - when using spectrophotometric methods[3]. Bias may also arise due to the intrinsic nuances of the method - for example, non-compliance with time intervals, errors in seeding the required number of cells, or the effect of the dye itself[4] [5].

In this paper, we propose a new method that would minimize operator influence and maximize convergence at least within a single method. The described method is mainly suitable for mass high-throughput screening. The principle of the method is based on the use of modified cell lines expressing fluorescent protein (this can be GFP, RFP, BFP, etc.). If a cell dies, the fluorescent protein is released into medium (similar to the LDH enzyme in the test of the same name) and can be detected by spectrofluorimeter.

General principle of the proposed method and its workflow:

1. Generation of a cell line stably expressing a fluorescent protein. For method reproducibility and stable expression, we recommend the use of lentiviral vectors.
2. Seeding of cells into wells of the plate. The first experiments described in the sections below were performed on a 96-well plate, however, for convenience, 48-well plates can also be used.
3. Addition of test compounds to study their suspected cytotoxicity. After the required period of time - lysis of cells in the wells designated as a negative viability control with 0.5-1% Triton X100 (or other detergent of choice).
4. Centrifugation of the plates for sedimentation of debris and detached cells.
5. Sampling a growth medium for fluorimetry.

Schematically the principle of the method is shown in Figure 1.

**Figure 1a.**
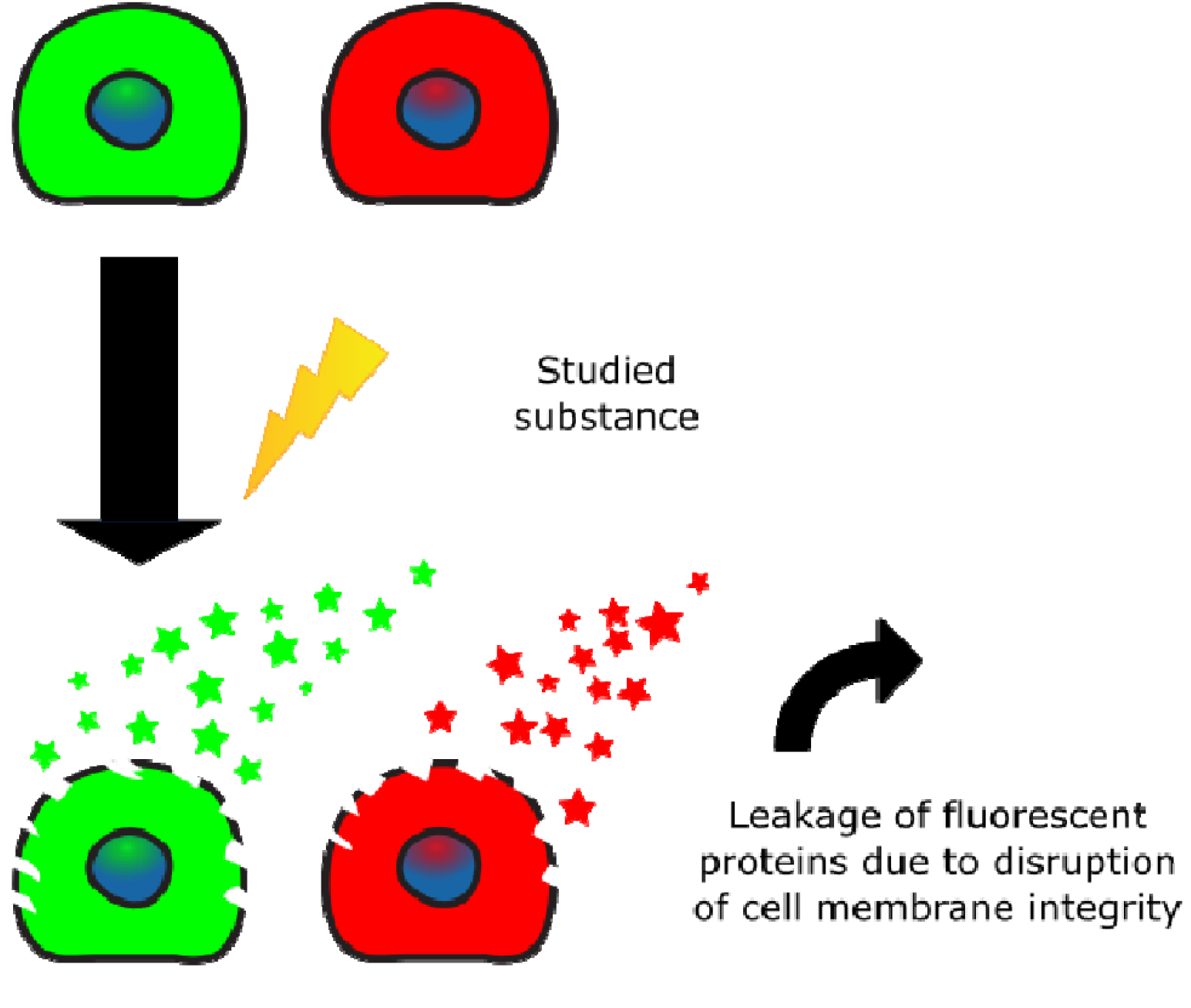
General principle of the proposed method.

**Figure 1b.**
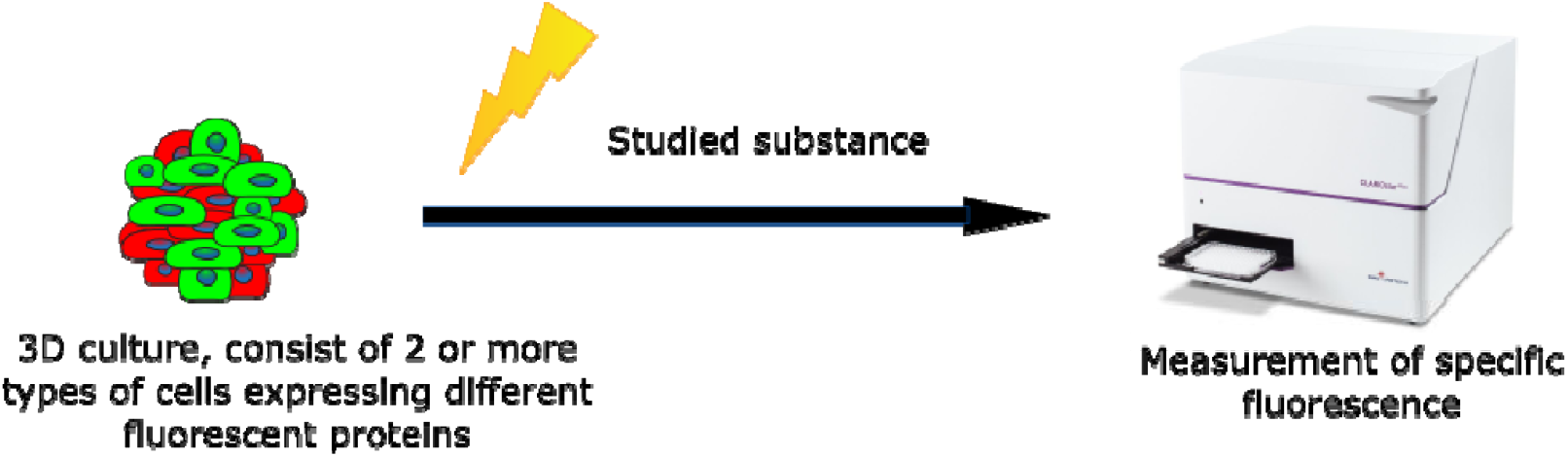
Proposed application of the method to 3D cultures. Advantages and disadvantages of the method –

*Advantages*

1. The method does not require chemical reagents other than TritonX100.
2. The method is not very sensitive to handling and accuracy of the operator.
3. The method can be applied to 3D-cultures and co-cultures. With the correct choice of excitation and emission wavelengths, it is possible to detect up to 2-3 signal bands from unmodified fluorescent proteins. The capabilities and palette of the method can theoretically be expanded by using narrow-band engineered fluorescent proteins or developing strategies to compensate the fluorescence spillover.
4. The method is easily scalable and can be applied to high-throughput screening in drug development.

*Disadvantages*

1. The method has little justifiable applicability outside of high-throughput screening, because generating each stable cell line is a rather labor-intensive process. To mitigate this drawback, we propose a solution in the form of mass generation of xFP-expressing cell lines from the NCI60 list[6].
2. To successfully perform this method, there are certain requirements for the formulation of the growth medium -namely the absence of phenol red[7]. Otherwise the method has a low signal-to-noise ratio. In addition, this method has limited applicability if the substances of interest have fluorescence properties.

### Selection of drugs and cell lines for experiments

We selected cell lines Huh7 and NCI-H1299 for these experiments.

Huh7 line, established in 1982 from a well differentiated hepatocyte derived cellular carcinoma cell line that was originally taken from a liver tumor in a 57-year-old Japanese male[8]. This cell line retained many phenotypic features of the original tissue, namely, the expression of a wide range of cytochromes responsible for the metabolism (activation, conversion, detoxification) of xenobiotics[9]. This property is especially important in in-vitro toxicological studies and translational research[10]. Cell line NCI-H1299 was established from a lymph node metastasis of the lung from a patient who had received prior radiation therapy[11]. This cell line is one of the most common in lung cancer research[12] [13] [14], radiotherapy[15] [16] [17] and chemotherapy[18] [19] [20].

For the experiments described in this paper, we chose capecitabine as an example, because this drug is mostly metabolized (activated) by liver cells.

### Experimental section

#### Materials and methods

As a basal growth medium we used phenol red-free F12 (PanEco, S600p). A basal growth medium was supplemented with fetal bovine serum (FBS) (HiMedia, RM10971-500ML), penicillin-streptomycin (PanEco, A073p’) and L-glutamine (PanEco, F032). As dissociating agents during passaging, we used a versene solution (PanEco, P080p) and trypsin-EDTA (PanEco, P039p).

#### Generation of the modified cell lines

In this paper, we used cell lines Huh7, NCI-H1299 and their derivatives, which were created by stable transduction with lentiviral vectors carrying the Tag-GFP2 (Evrogen, LP004[21]) and Tag-RFP (Evrogen, LP001[22] [23]) genes[24].

Thus, after the stages of cloning and selection, performed according to standard methods[25], we obtained the following cell lines - Huh7-GFP, NCI-H1299-GFP, NCI-H1299-RFP.

For the sake of reproducibility, we strongly advise against using lines obtained by simple transfection with a non-integrating vector, because the presence of selecting antibiotics (e.g. G418) may interfere with viability measurements.

#### Design of roof-of-concept experiment

The described experiment serves to establish the functionality of the principle of the method and answer the questions -

1. Is it possible to detect a fluorescent signal from a fluorescent proteins that were released as a result of cell death or lysis?
2. Is 2-channel detection of independent of such fluorescent signals possible?

The experiment involved 2 cell lines, derivatives of NCI-H1299 – NCI-H1299-GFP and NCI-H1299-RFP, expressing GFP and RFP respectably.

The cells were seeded into the wells of a 96-well plate in the amount of 20 thousand cells per well. The cells were seeded in 6 rows of wells - 2 rows for a line expressing green fluorescent protein, 2 rows for a line expressing red fluorescent protein, and 2 rows for a mixture of cells from both lines. Of all the groups of cell lines, one row was for the experiment, the other for the control.

After 24 hours of incubation in a humidified CO2 incubator at 37*C and 5% CO2, 10 μl of medium was added to the control wells, and 10 μl of 10% Triton-X100 was added to the experimental wells. The contents of the wells, to which the detergent was added, then mixed by pipetting. Then the culture plate was placed in the incubator for 20 minutes to lyse the cells, then the contents of the wells with detergent were once again carefully mixed by pipetting. The plate was placed in a centrifuge with adapters for the plates and centrifuged at 1000 G for 10 minutes. 50 µl was taken from each well and transferred to a new plate. 50 μl of complete growth medium was added to three empty wells to use as a blank control. This new plate was then measured on the BMG Clariostar plate reader in fluorimetry mode.

Readings were performed on these wavelengths:

For RFP: excitation - 554 nm, emission – 585 nm.

For GFP: excitation - 488 nm, emission – 516 nm.

#### Experiment to determine the toxicity of capecitabine using described method

For this experiment, cell lines NCI-H1299-RFP and Huh7-GFP were taken. Capecitabine was chosen to demonstrate the capabilities of the method.

Cells were seeded at a concentration of 20,000/well of a 96-well plate. The following combinations were used: NCI-H1299-RFP monoculture, Huh7-GFP monoculture, NCI-H1299-RFP/Huh7-GFP coculture, and NCI-H1299-RFP/Huh7 coculture. For cocultures, 10,000 cells of each line were seeded per well. The experiment was carried out in triplicate. For each mono- and coculture, their own wells were allocated for intact control (positive viability control) and negative viability control (at the end of the experiment, the cells in these wells will be lysed). After 24 hours, the growth medium in the wells was replaced with a medium containing the studied compound (capecitabine), in the wells for controls a simple replacement of the medium was performed.

After 48 hours of incubation in a humidified CO2 incubator at 37*C and 5% CO2, 10 μl of medium was added to the control wells, and 10 μl 1% Triton X100 was added to the experimental wells. The contents of the wells to which the detergent was added were mixed by pipetting. Then the culture plate was placed in the incubator for 20 minutes to lyse the cells, then the contents of the wells with detergent were once again carefully mixed by pipetting. The plate was placed in a centrifuge with adapters for the plates and centrifuged at 1000 G for 10 minutes. 50 µl was taken from each well and transferred to a new plate. 50 μl of complete growth medium was added to three empty wells to use as a blank control. This new plate was then measured on the BMG Clariostar plate reader in fluorimetry mode.

### Experiment to determine the applicability of the method to 3D cultures (spheroids and alginate hydrogel)

#### Spheroids

For this experiment, cell lines NCI-H1299-RFP and Huh7-GFP were taken. The spheroids were obtained by the hanging drop method. Briefly, rows of drops with a volume of 25 μl were placed on the inside of the top lid of a Petri dish, each drop containing 2000 cells. 10 ml of PBS was then added to the bottom of the Petri dish and the top lid was carefully inverted and placed on top. After 5 days, the spheroids were formed.

The medium was then changed twice, 10 μL at a time. Then after, 5^th^ day, 10 μl of medium in droplets, which contained spheroids for the experiment, was replaced by 10 μl of complete culture medium containing 375 μM capecitabine. Thus, the final concentration of capecitabine in 25 μl was 150 μM. This concentration was chosen based on the results of a previous experiment.

After 48 hours of incubation in a humidified CO2 incubator at 37^*^C and 5% CO2, 10 μl of medium was taken from each group of droplets and pooled into their respective wells. The entire contents of the drops designated as “negative viability control” were taken along with the spheroids and transferred into a 250 μl PCR tube. 6 µl of Triton-X100 was added there. The tube was then vortexed and left to incubate on a rotator at room temperature for half an hour before being mixed again and centrifuged at 1000g for 10 minutes to pellet debris. 60 µl from this tube was transferred to the corresponding well of the plate. This new plate was then measured on the BMG Clariostar plate reader in fluorimetry mode.

#### Alginate hydrogel

For the purposes of this work, an experiment was conducted to create three-dimensional “droplets” of cross-linked sodium alginate with cells enclosed inside. The protocol outlined in the work[26] with minor modifications was taken as a basis. The volume of spheroids was changed - 100 μl, the order of addition and composition of the cross-linking solution CaCl2, the concentration of sodium alginate and the number of cells were changed. The sodium alginate concentration of 2.5% stated in (Smit et al., 2020) turned out to be irreproducible in practice - uniform stirring and manipulation of such a solution turned out to be impossible due to its excessive viscosity. After a series of preliminary experiments, it was decided to settle on a concentration of 1.1%, and use 1% CaCl2 in F12 as a crosslinking solution.

Before starting the experiment, a solution of 1.1% sodium alginate in Hanks’ buffer was prepared. 1.1 grams of sodium alginate was added to 100 ml of Hanks’ solution in a heat-resistant flask. Stirring was carried out with a magnetic armature for 4 hours at a temperature of 55C. After stirring all visible lumps and inhomogeneities, the temperature on the magnetic stirrer was raised to 200C and the solution was brought to a boil twice for sterilization (since its sterilization through filters is impossible due to its viscosity). After which the flask with the solution was introduced into the laminar flow and the solution was aseptically transferred into a test tube for cooling.

A solution for “crosslinking” was also prepared - F12 with the addition of sterile calcium chloride to a concentration of 1%. To perform this experiment, cell lines H1299-RFP, H1299-GFP, Huh7-GFP and Huh7 were taken. Cell cultivation was carried out according to standard procedures. Upon reaching 80-90% confluency, Petri dishes with cultures were washed with versene, the cells were trypsinized with a 0.25% trypsin-EDTA solution, and the cell suspension was transferred to centrifuge tubes. After centrifugation at 1000 g for 5 minutes, the supernatant was removed and the cells were resuspended in 2 ml of fresh F12 medium. The concentration of the suspensions in all cell lines was adjusted to 1 million/ml, and 1 ml of the suspension of each line was transferred into a 1.5 ml Eppendorf, in the case of cocultures - 0.5 ml of each.

Thus, the following were prepared for the experiment:

1. Monocultures – H1299-GFP, H1299-RFP, Huh7-GFP
2. Cocultures – H1299-GFP/Huh7, H1299-RFP/Huh7 and H1299/Huh7-GFP.

After pippeting, tubes with cell suspensions were centrifuged at 1000g for 10 minutes. In parallel, two 12-well plates were taken, and 4 ml of cross-linking solution was added to each well. After centrifugation, the supernatant was removed and 1 ml of Hanks’ solution with sodium alginate was added to each tube.

After slow and careful mixing, 2 drops of a cell suspension in sodium alginate were dripped into the wells of the plate from a height of 5-7 cm. The volume of each drop is 100 µl. This height is optimal for such viscous droplets that they acquire a spherical shape upon contact with the cross-linking solution. The excavation was carried out according to the following scheme indicated in the Table 1:

**Table 1.** Scheme of sodium alginate droplets.

|  |  |
| --- | --- |
|  | Plate №1 |

| Cell line | Capecitabine | Negative viability control | Positive viability control |
| --- | --- | --- | --- |
| H1299-GFP | 2 droplets | 2 droplets | 2 droplets |
| H1299-RFP | 2 droplets | 2 droplets | 2 droplets |
| Huh7-GFP | 2 droplets | 2 droplets | 2 droplets |
| Plate №2 |  |  |  |
| H1299-RFP/Huh7-GFP | 2 droplets | 2 droplets | 2 droplets |
| H1299-RFP/Huh7 | 2 droplets | 2 droplets | 2 droplets |
| H1299/Huh7 | 2 droplets | 2 droplets | 2 droplets |

The crosslinking solution was then removed from all wells, the wells were washed twice with F12, and 2 ml of fresh F12 was added to each well. 3D cultures in alginate drops were cultivated for 14 days, the medium was replaced every 2 days. As visible three-dimensional structures grew and formed in the alginate droplets (shown in the figures), the formation of spheroids was monitored (Figure 2).

**Figure 2a.**
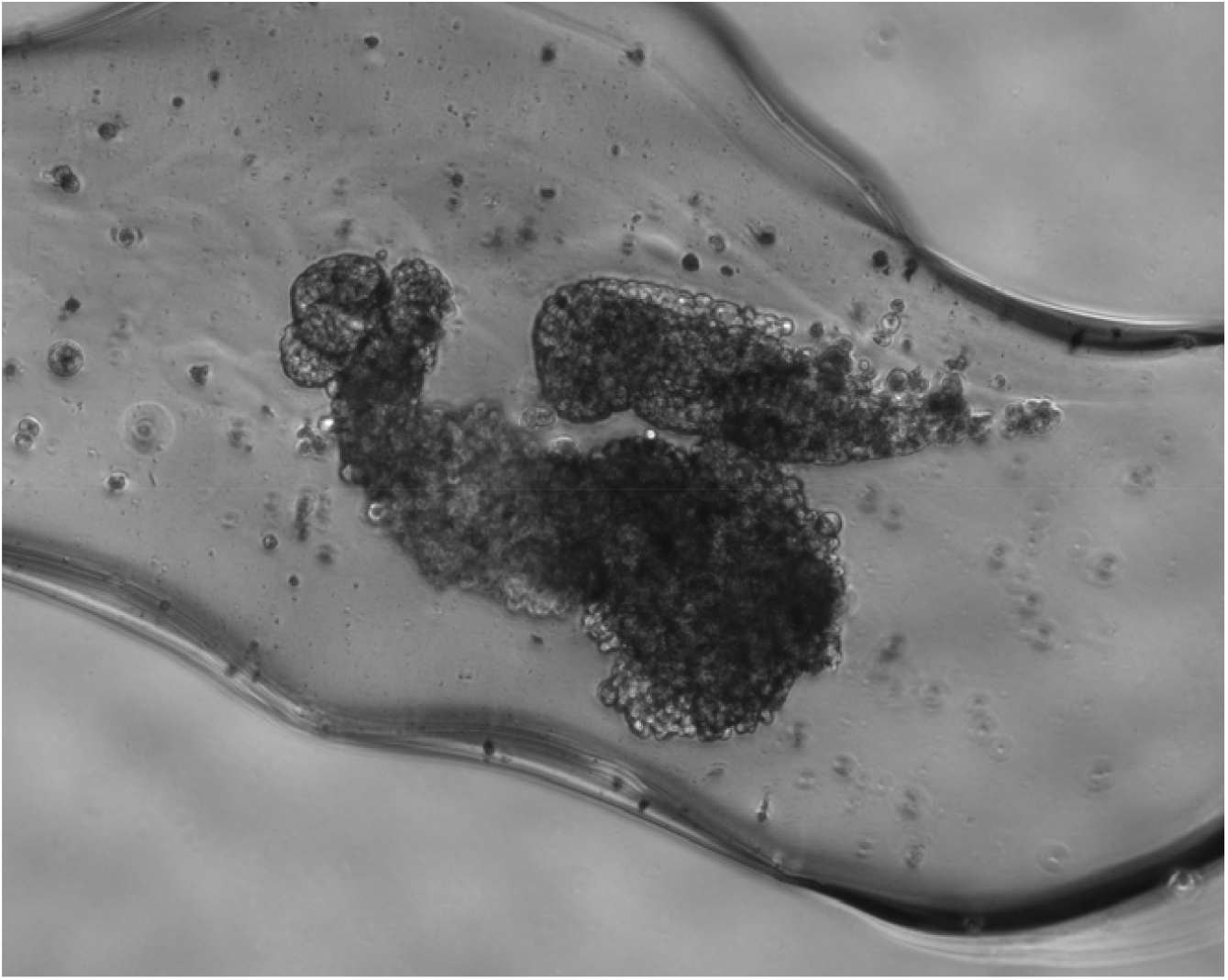
H1299-GFP culture growth, phase contrast microscopy. 5th day of cultivation.

**Figure 2b.**
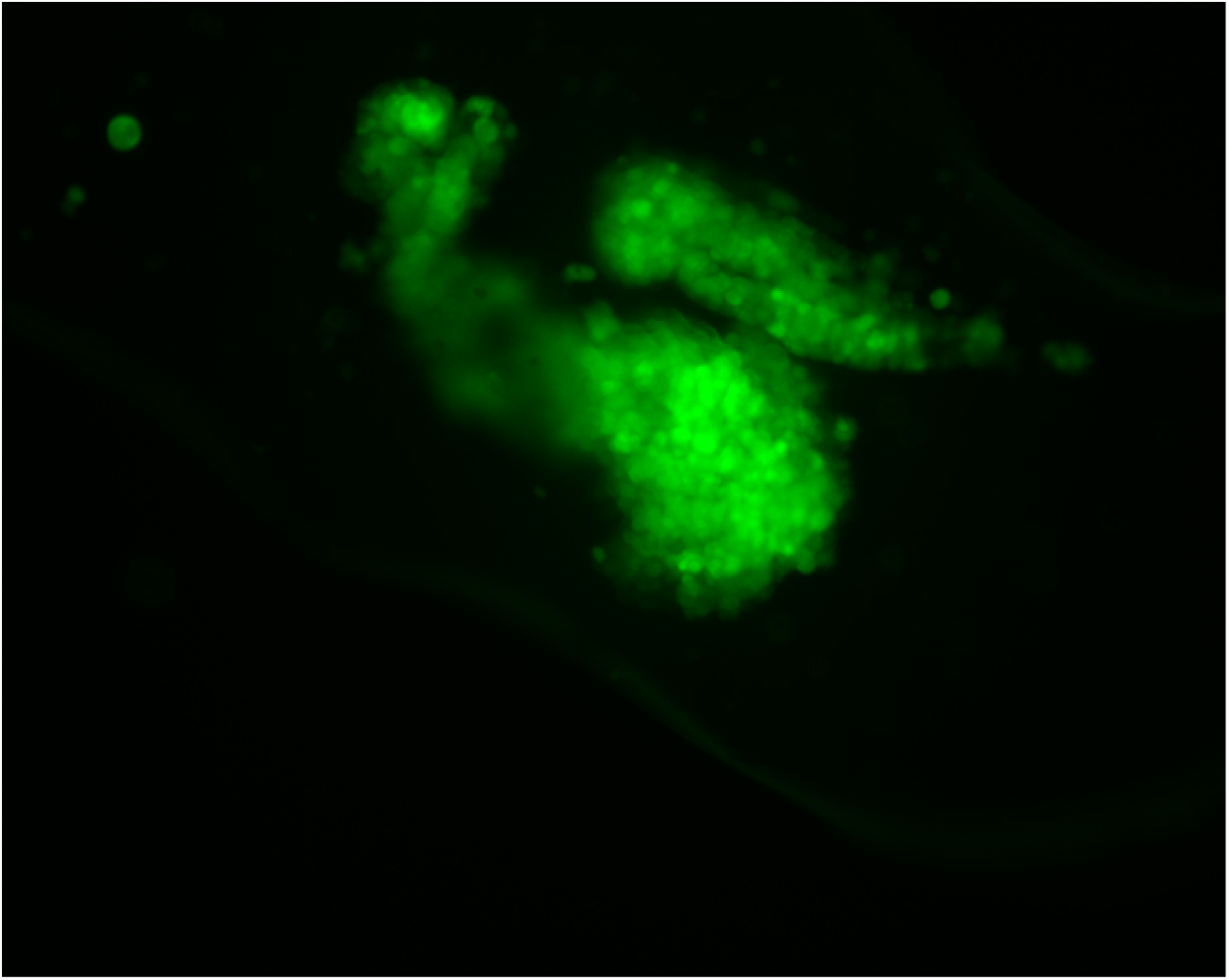
Growth of the H1299-GFP culture, fluorescence in the green channel. 5th day of cultivation.

Interestingly, the Huh7 and Huh7-GFP cultures, when growing in an alginate matrix, along with spheroids (at the later stages of cultivation - days 9-12) began to form elongated, radially located formations (Figure 3).

**Figure 3.**
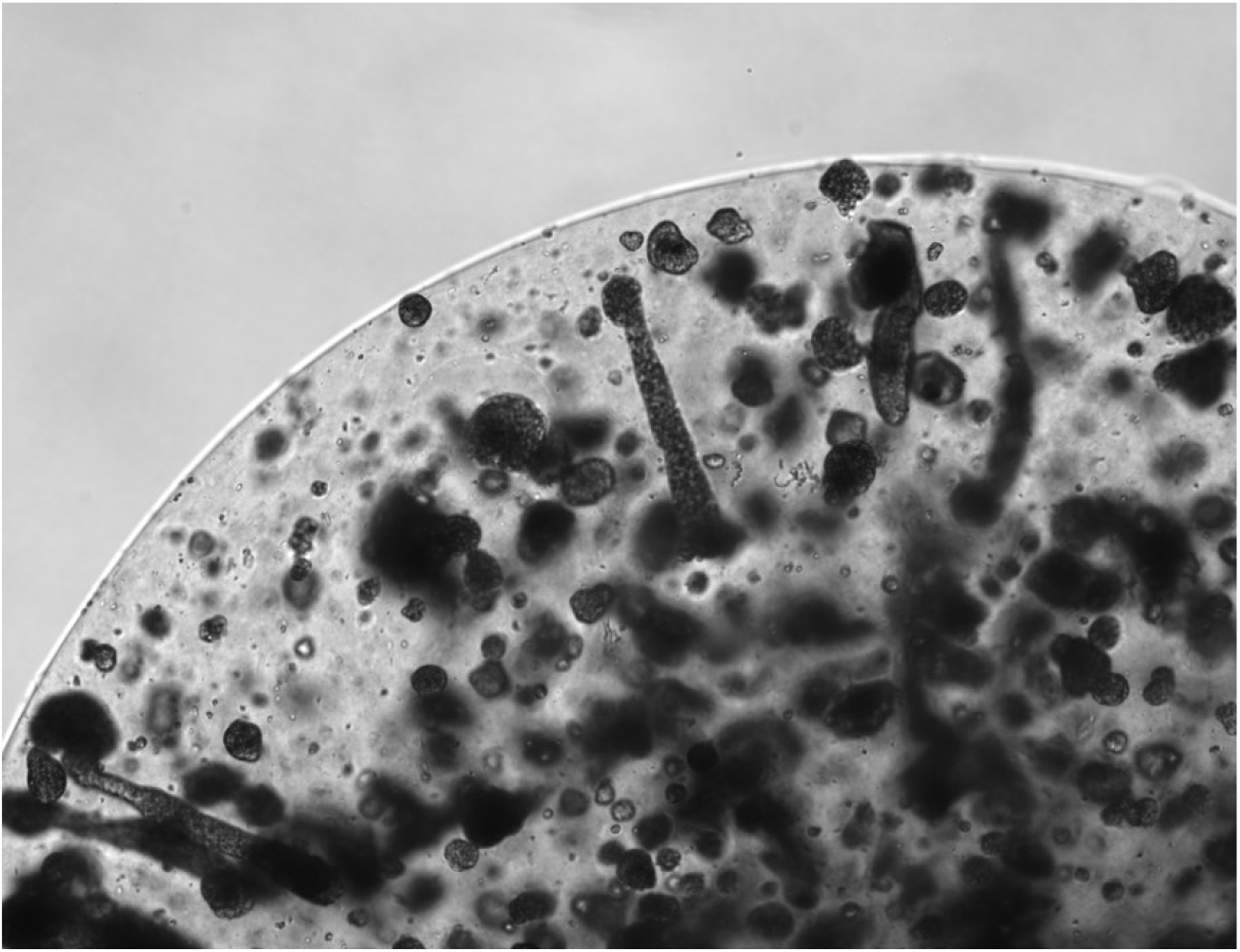
Huh7 culture, 11th day of cultivation, phase contrast microscopy.

On the 14th day of cultivation, the medium in the wells was replaced with a medium containing 150 μM capecitabine (for the corresponding wells). In control wells, the medium was replaced with regular F12 medium. After 36 hours, 20 μl of Triton X100 was added to the negative control wells. After another 12 hours, 100 μl of medium was taken from each well of each culture into the wells of a 96-well plate, and fluorescence was measured in the GFP and RFP channels on a CLARIOStar BMG spectrofluorimeter. Another day later, an additional measurement was performed. Thus, we got 2 time points - 48 and 72 hours.

## Results

### Proof-of-concept experiment

The results show great specificity of detection in the respective excitation and emission channels (Figure 4).

**Figure 4a.**
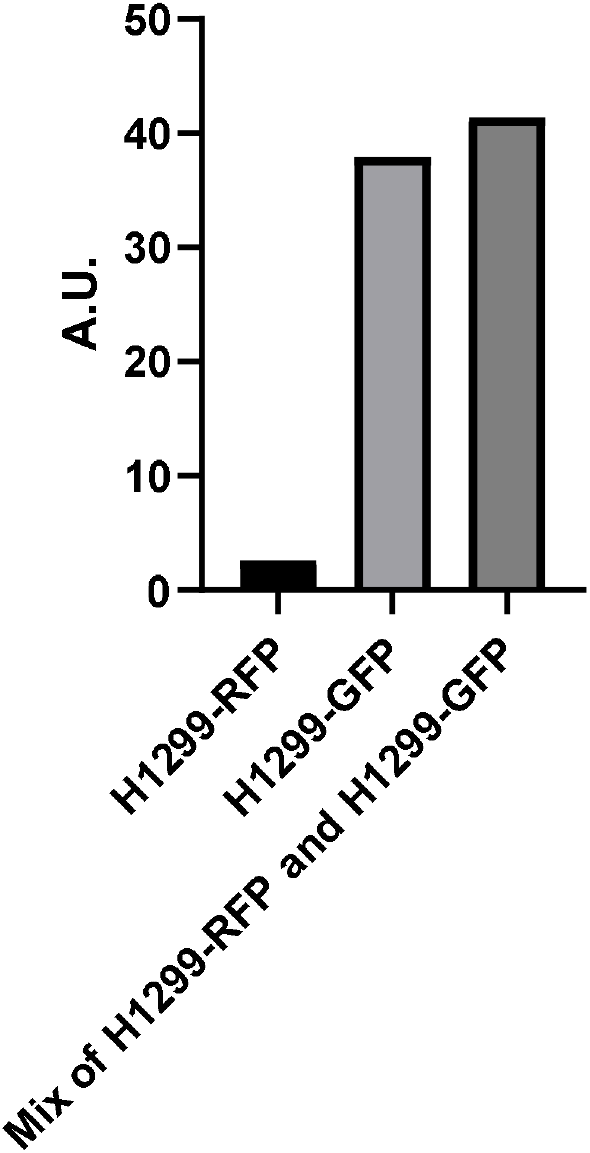
Specificity of cell lysate fluorescence detection in RFP channel.

**Figure 4b.**
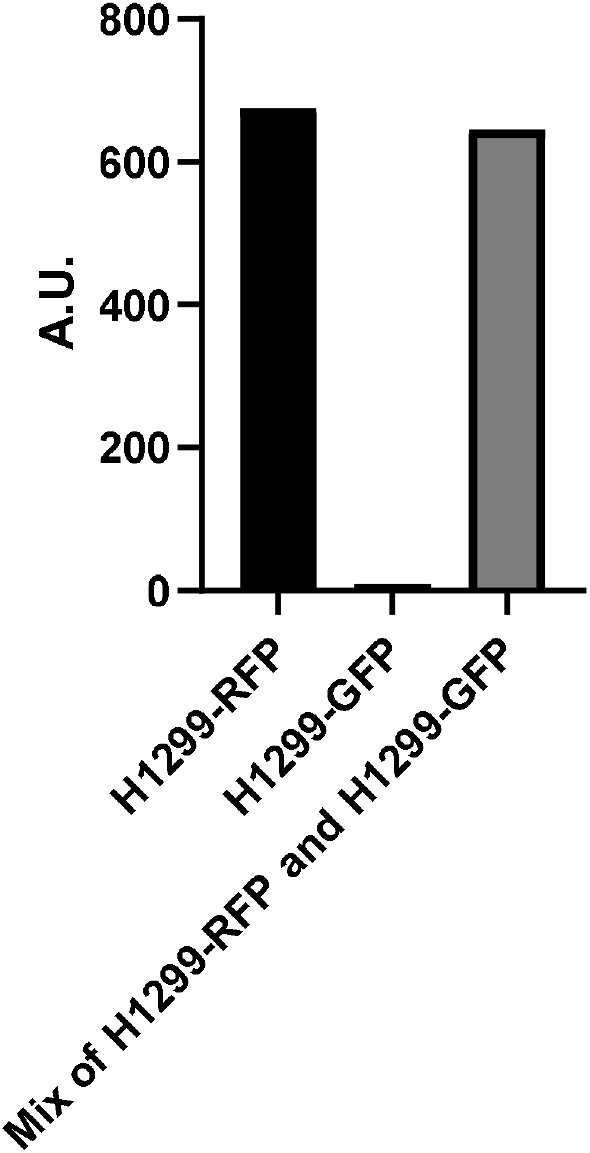
Specificity of cell lysate fluorescence detection in GFP channel.

### Determination of capecitabine cytotoxicity (2D cultures)

After measuring the fluorescence level, the results were calculated using the formula to determine viability:

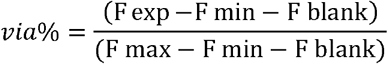

Where:

F blank – fluorescence of fresh medium

F exp – fluorescence of a medium from wells with previously added capecitabine

F min – fluorescence of a medium from wells with intact cells (positive control of viability)

F max – fluorescence of a medium from wells with lysed cells (negative control of viability)

Results from this formula can be expressed as any decimal fraction between 0 and 1 and then converted to a percentage. Where 0 is the absence of cell death, and 1 is - 100%, a full cell death.

Using this formula, results were obtained for the toxicity of capecitabine in mono- and co-cultures against NCI-H1299-RFP, NCI-H1299-GFP and Huh7-GFP cells (Figure 5). Note that the calculated TC50 (toxic concentration), in case of NCI-H1299-RFP and NCI-H1299-GFP, differs between RFP and GFP channels, possibly due to cell decomposition products which have a great fluorescence in GFP-like range and thus interfering with a results.

**Figure 5a.**
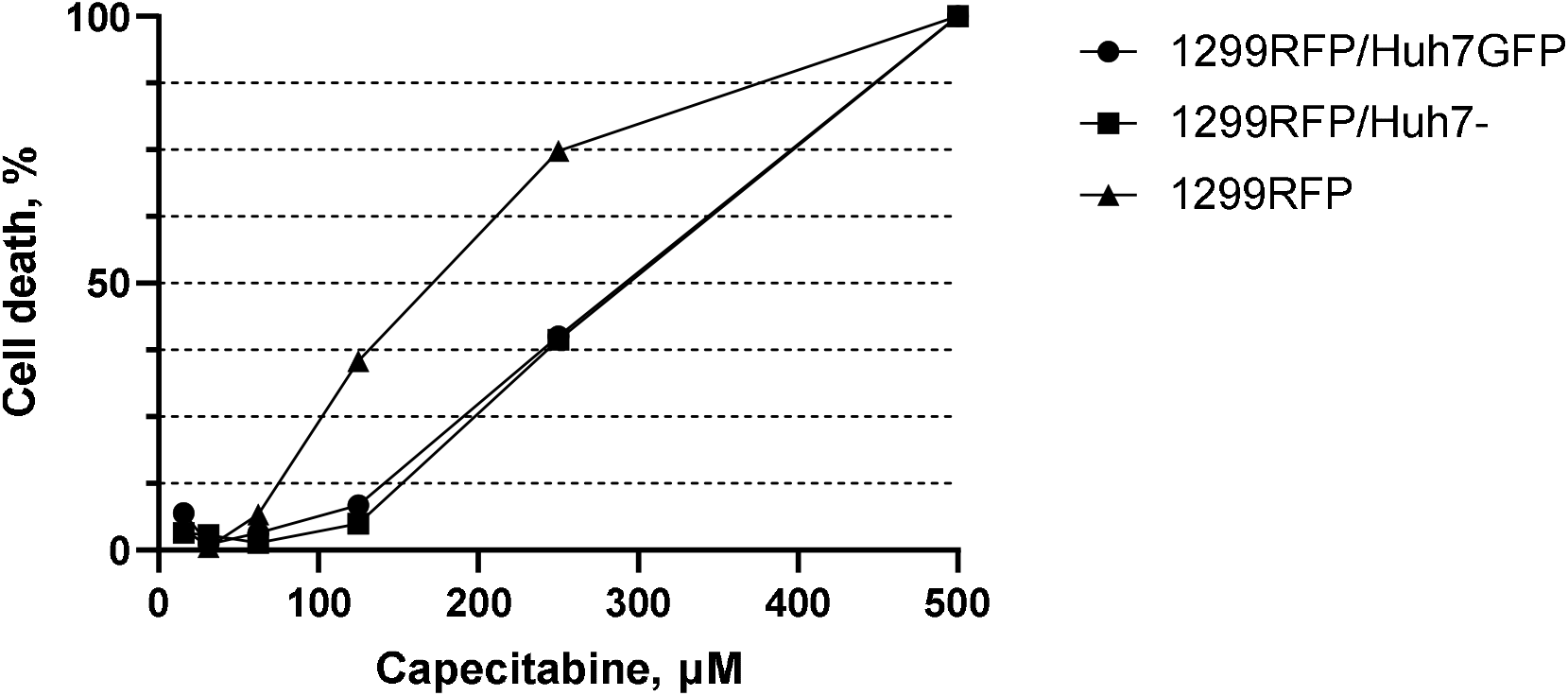
Viability of NCI-H1299-RFP in cases of mono- and cocultures.

**Figure 5b.**
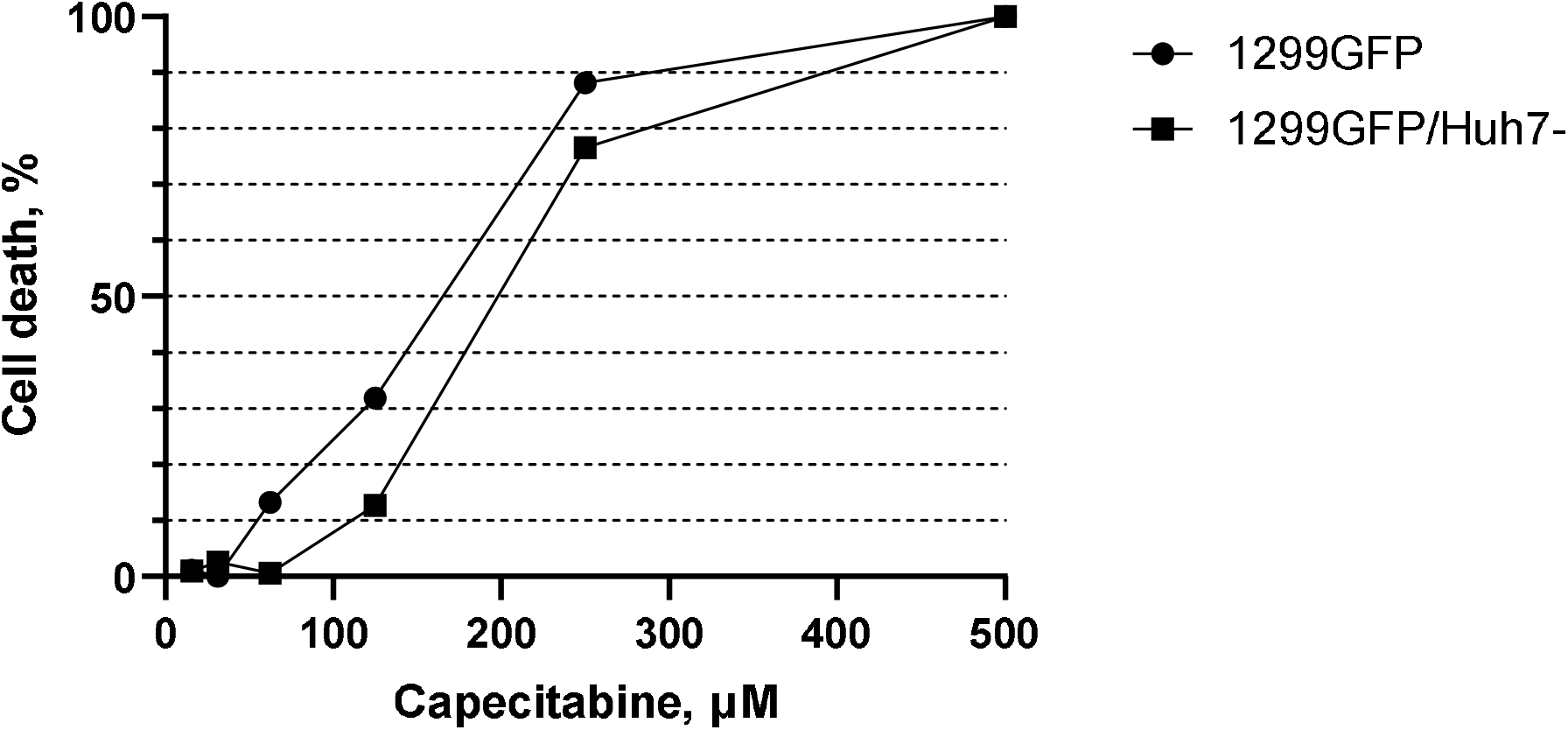
Viability of NCI-H1299-GFP in cases of mono- and cocultures.

**Figure 5c.**
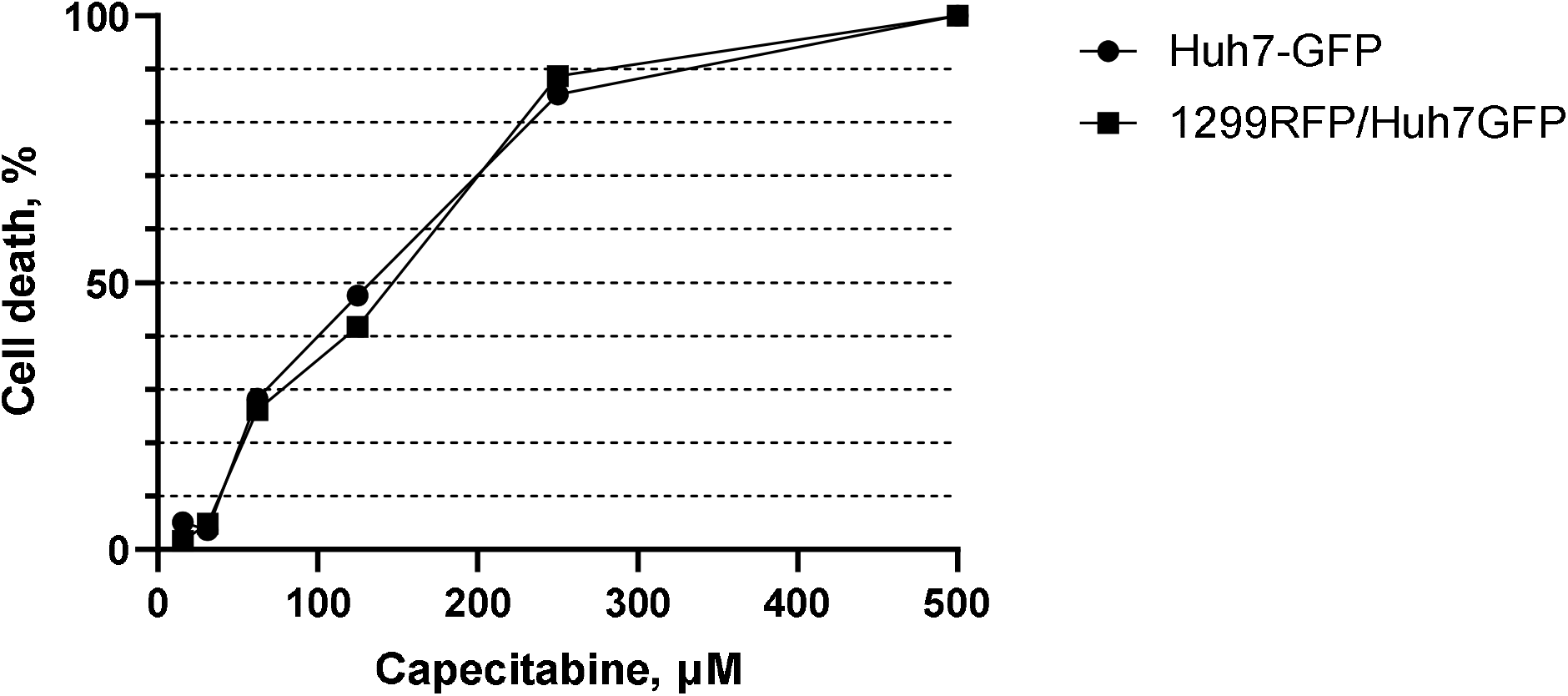
Viability of Huh7-GFP in cases of mono- and cocultures. Calculated IC50 for capecitabine:

Calculated IC50 for capecitabine:

NCI-H1299-RFP – 175,6 μM

NCI-H1299-RFP (coculture with Huh7-GFP) – 309,9 μM

NCI-H1299-RFP (coculture with Huh7) – 282,4 μM

NCI-H1299-GFP – 158,2 μM

NCI-H1299-GFP (coculture with Huh7) – 197,2 μM

Huh7-GFP – 137,8 μM

Huh7-GFP (coculture with NCI-H1299-RFP) – 146,3 μM

### Determination of capecitabine cytotoxicity (3D cultures)

#### Spheroids

The results show a significant reduction in the toxicity of capecitabine in the presence of metabolizer cells (Huh7) against NCI-H1299 cells (Figure 6a and 6b). Note that the viability of Huh7 cells in monoculture and coculture does not change as much, which indicates that the presence of Huh7 influences NCI-H1299, and not vice versa (Figure 6c).

**Figure 6a.**
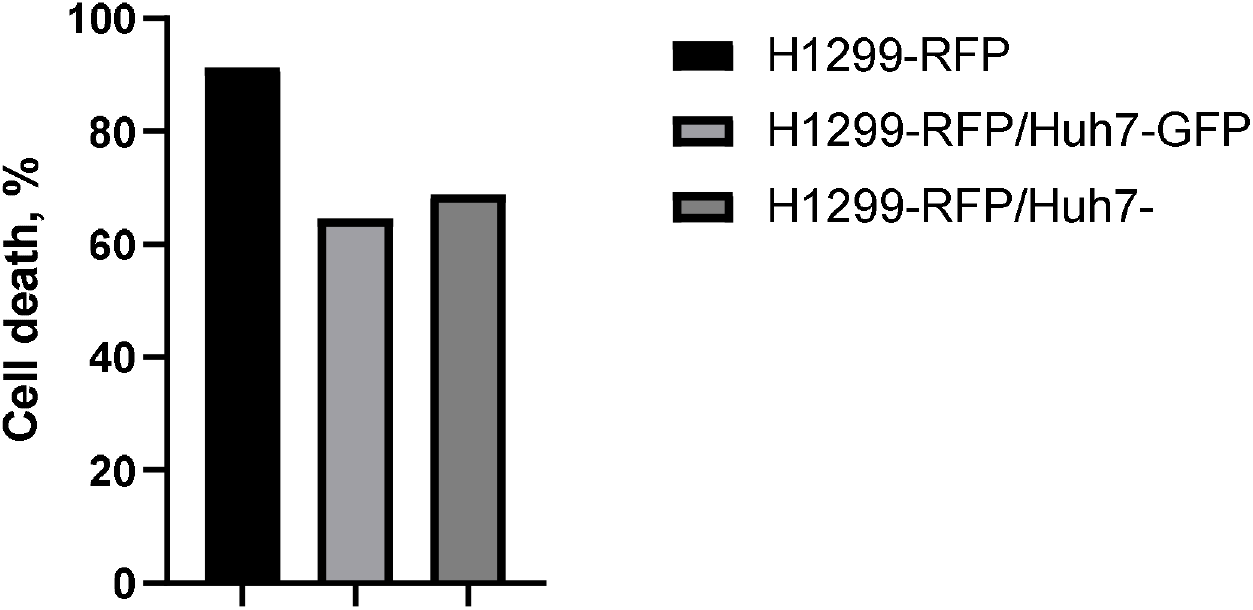
Viability of NCI-H1299-RFP cells in mono- and cocultures.

**Figure 6b.**
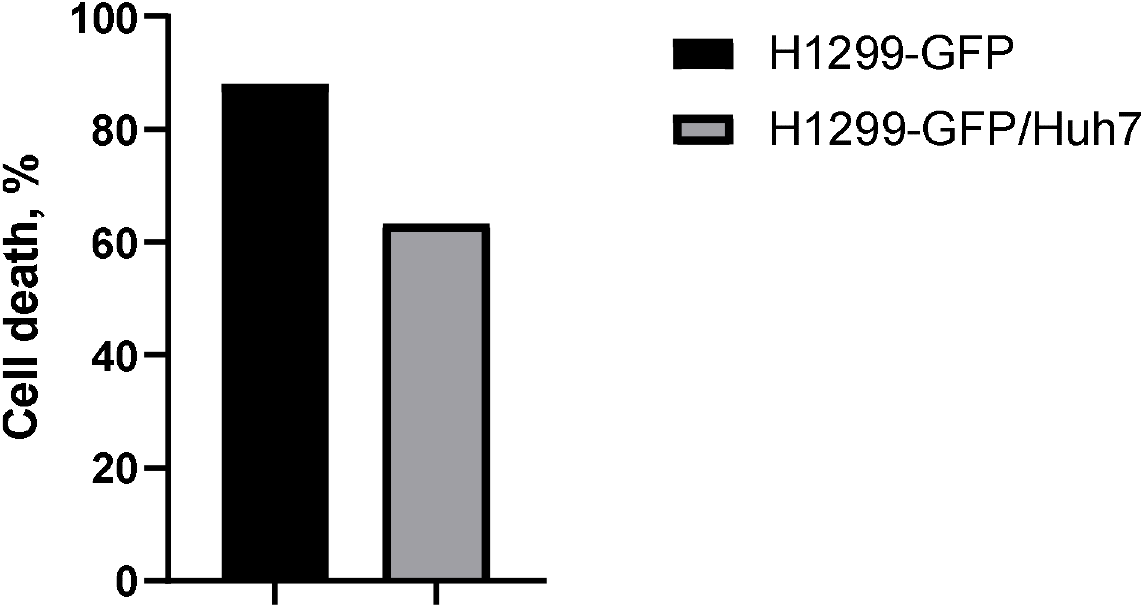
Viability of NCI-H1299-GFP cells in mono- and cocultures.

**Figure 6c.**
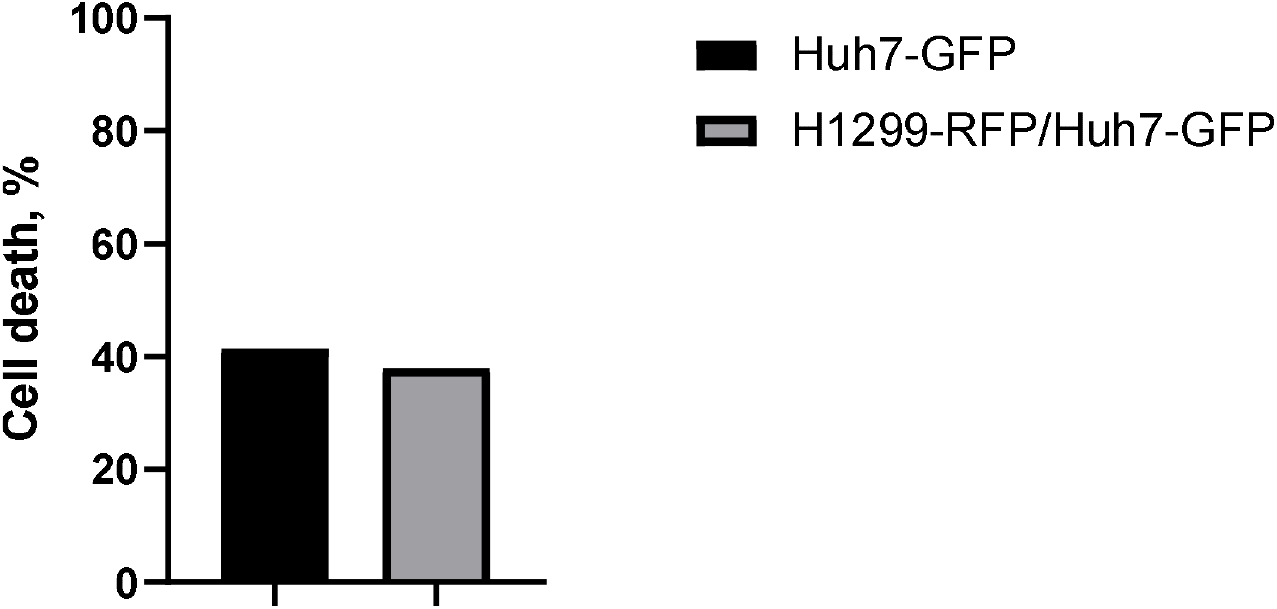
Viability of Huh7-GFP cells in mono- and cocultures

#### Alginate hydrogel

The results show a dramatic reduction in sensitivity to capecitabine. Interestingly, this happens in all cases (mono- and cocultures) (Figure 7). An additional experiment was carried out using the same method, where 300 µM capecitabine was taken into the experiment - sensitivity (i.e. cell death) increased, but not much (Figure 7 d, e, f).

**Figure 7a.**
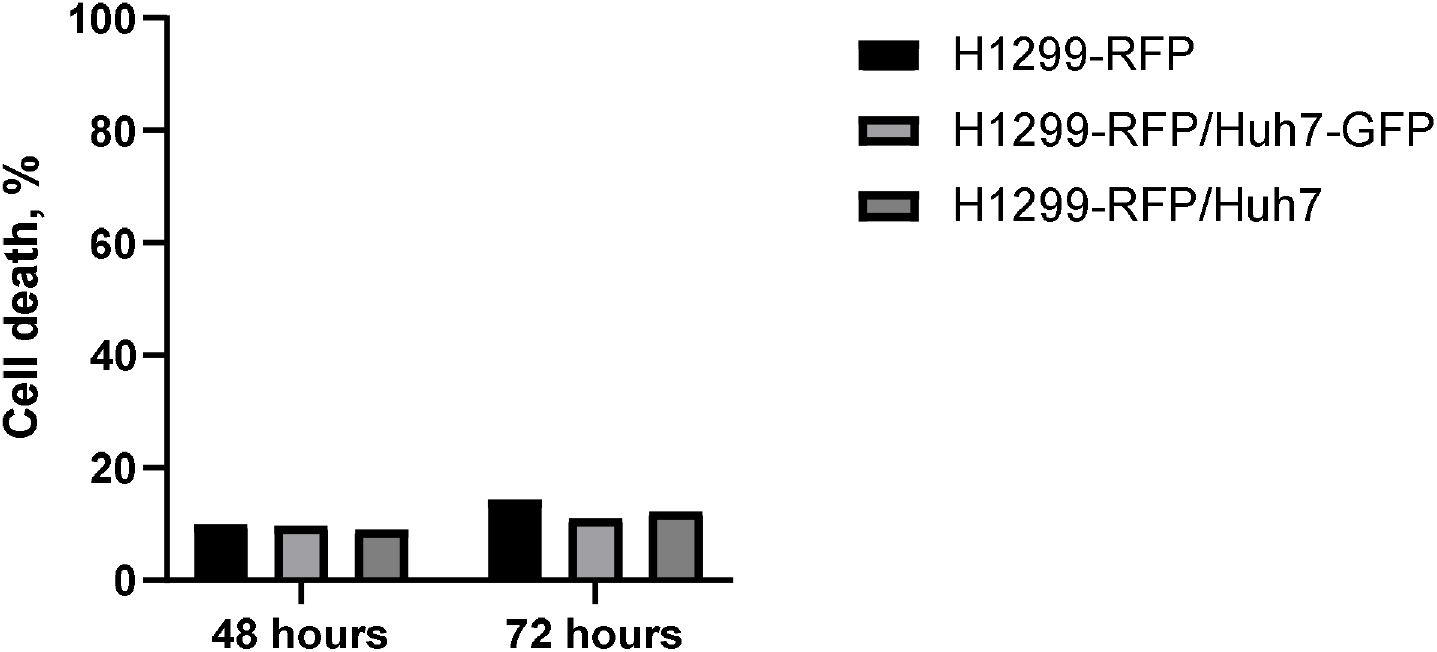
Viability of NCI-H1299-RFP cells in mono- and cocultures

**Figure 7b.**
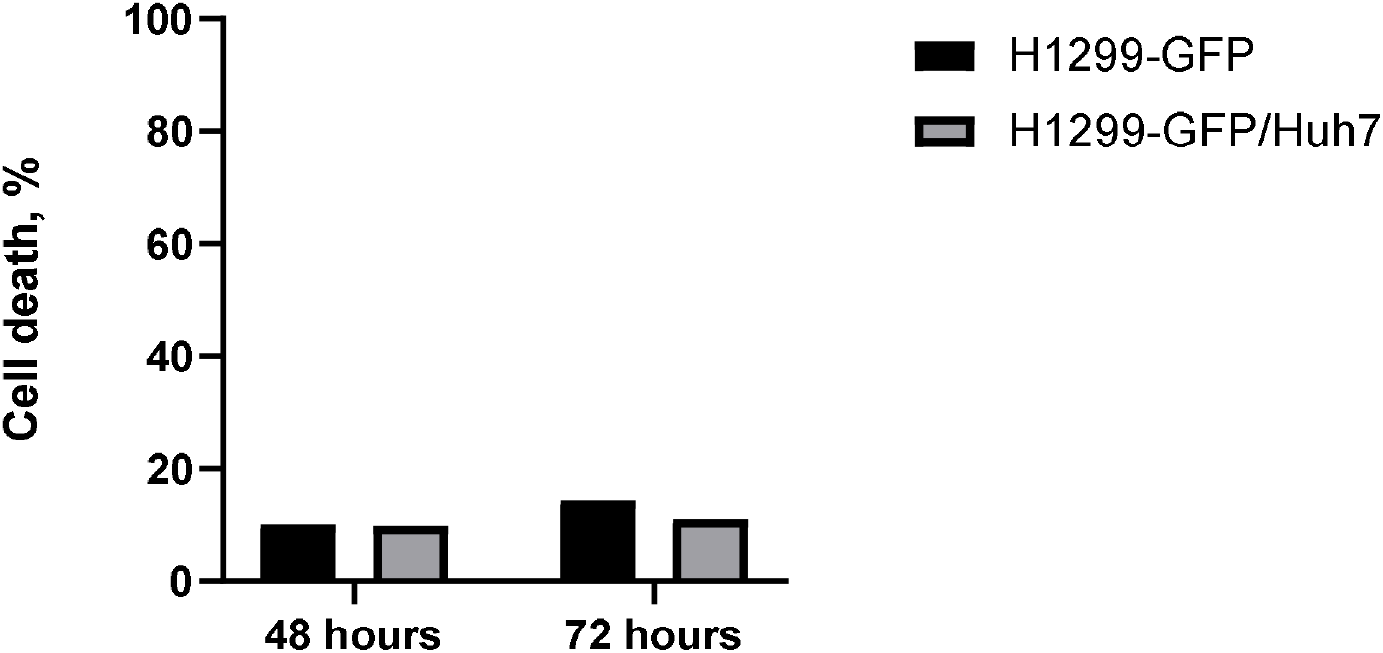
Viability of NCI-H1299-GFP cells in mono- and cocultures

**Figure 7c.**
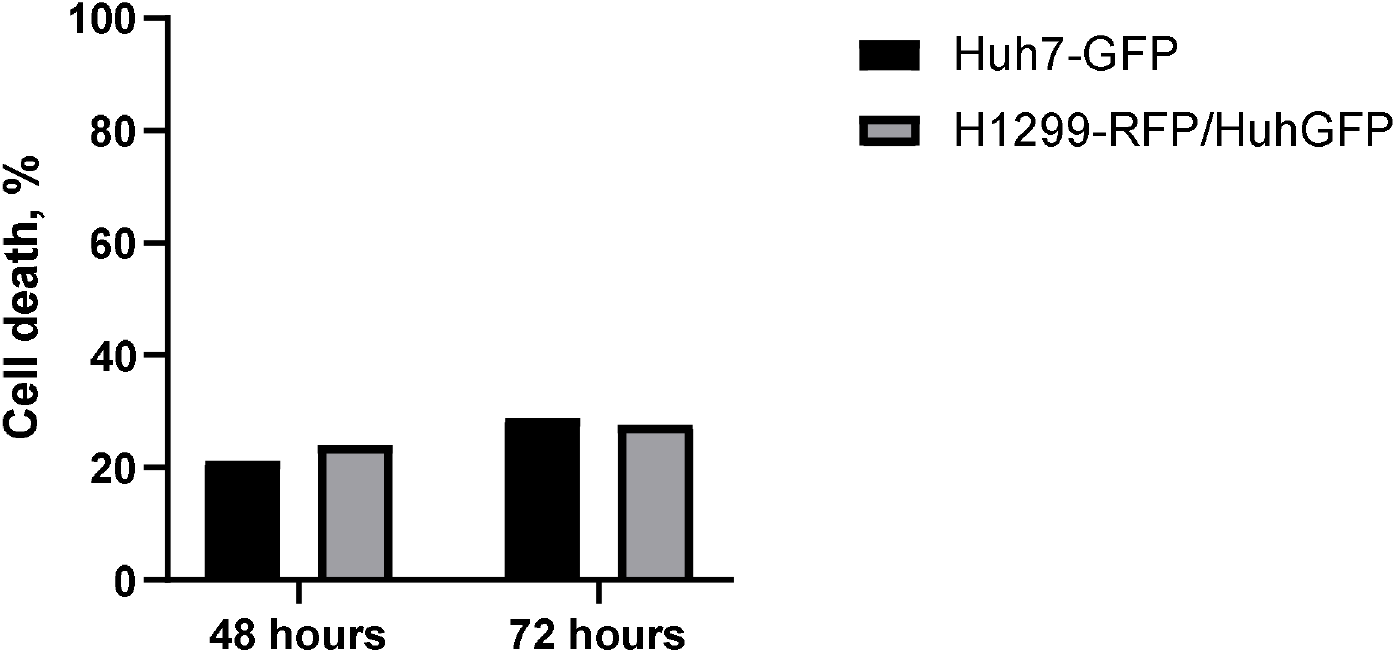
Viability of Huh7-GFP cells in mono- and cocultures

**Figure 7d.**
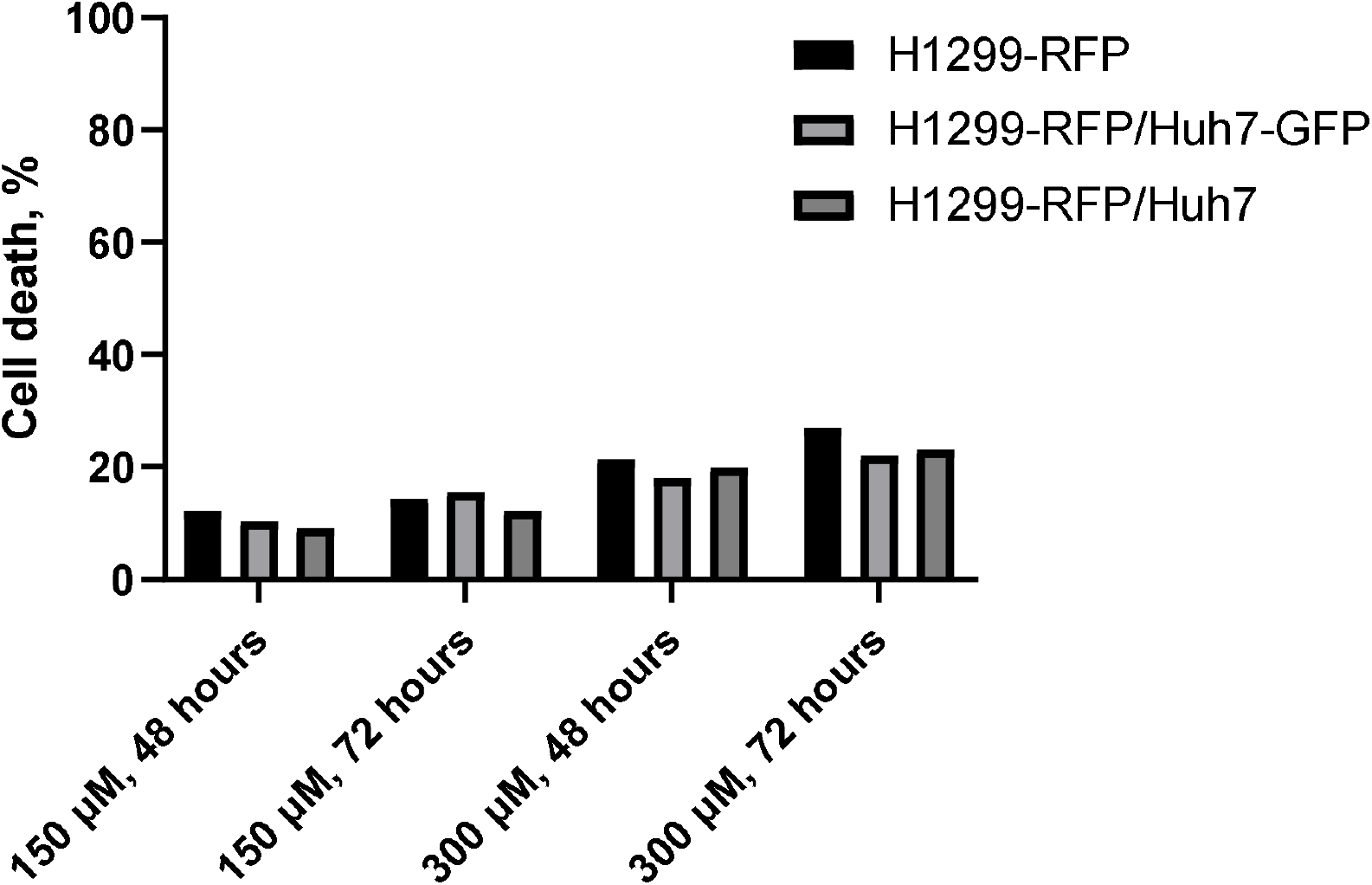
Comparison of NCI-H1299-RFP cells viability in the presence of 150 and 300 μM capecitabine.

**Figure 7e.**
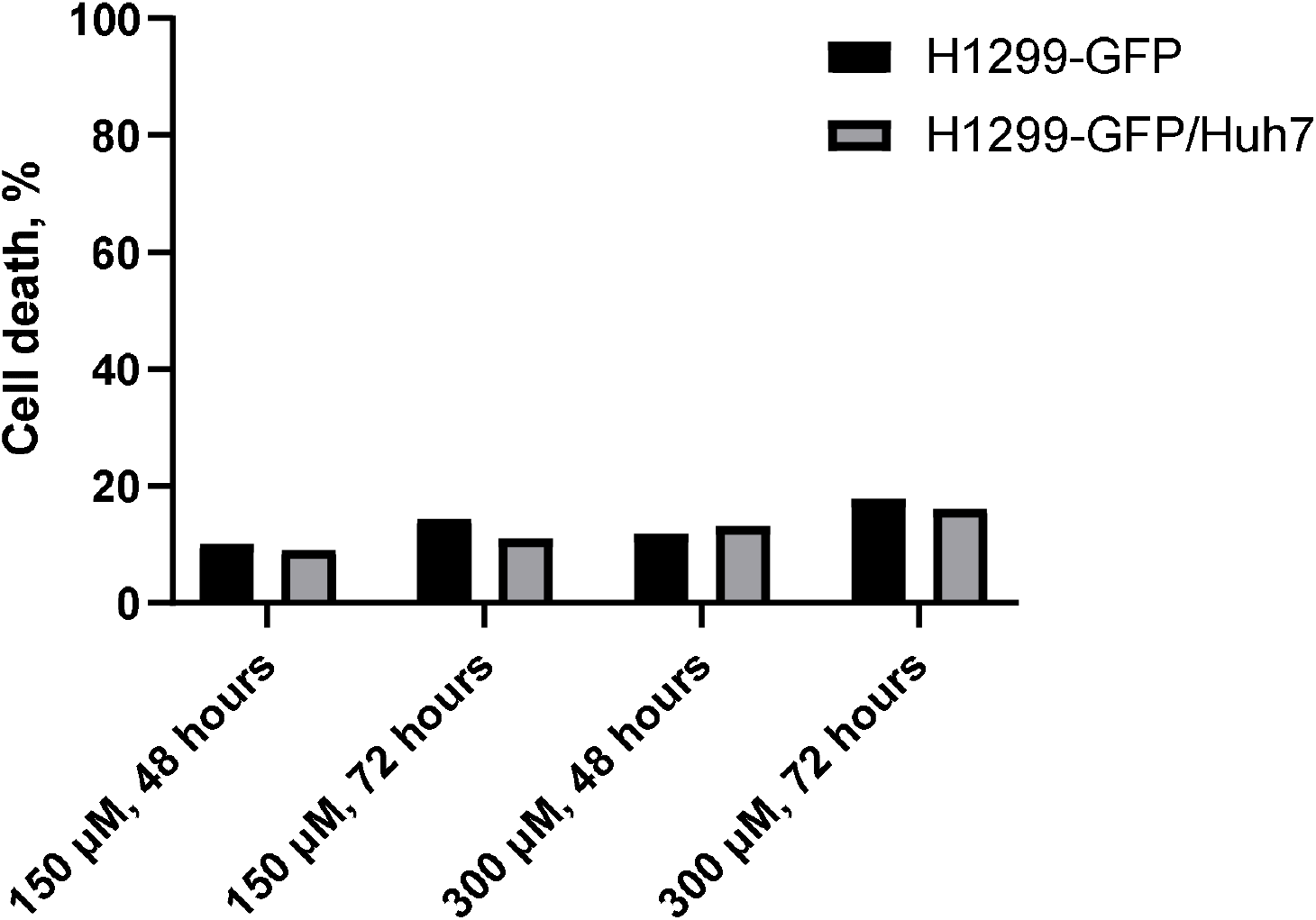
Comparison of NCI-H1299-GFP cells viability in the presence of 150 and 300 μM capecitabine.

**Figure 7f.**
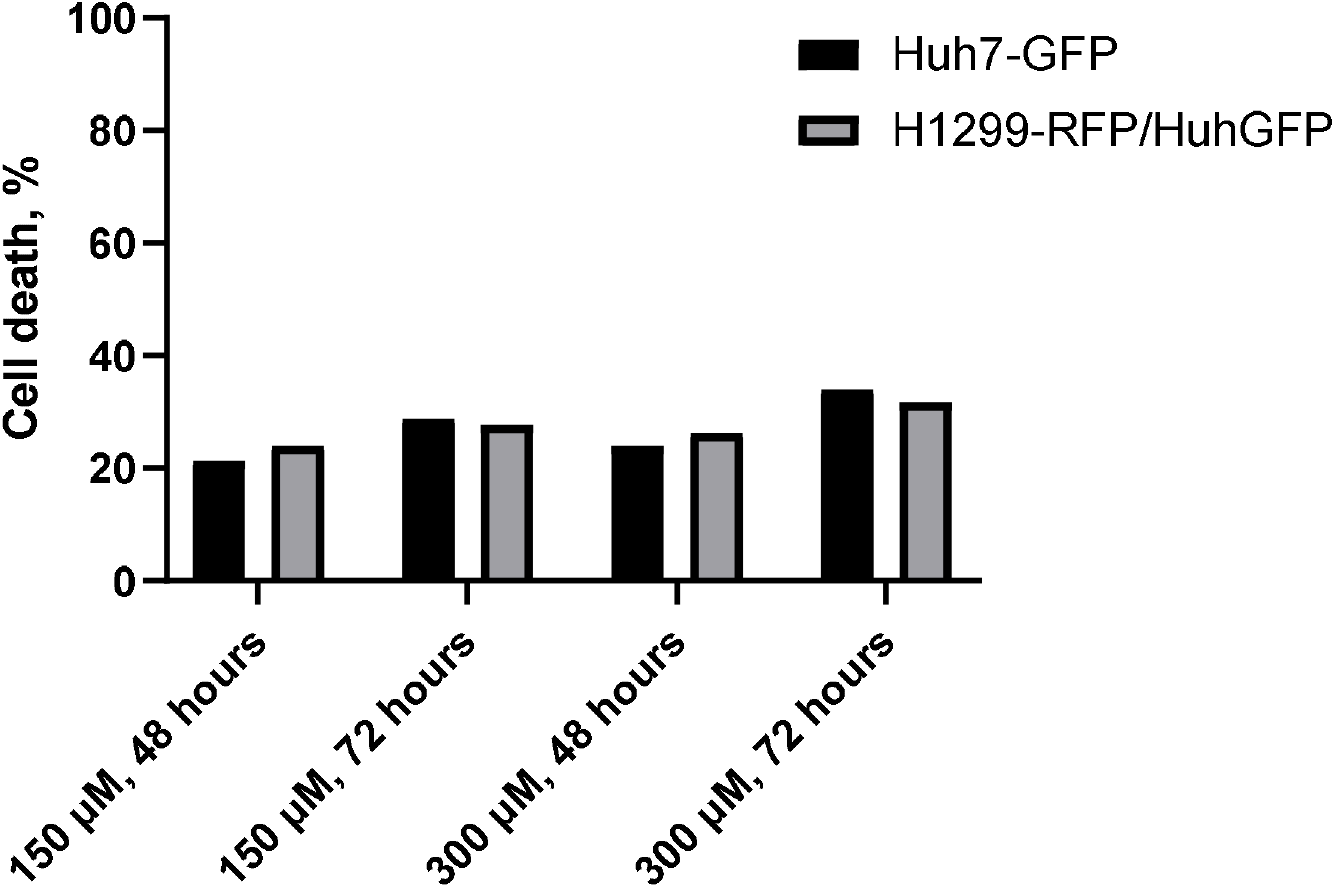
Comparison of Huh7-GFP cells viability in the presence of 150 and 300 μM capecitabine.

## Conclusion

This work was intended to demonstrate a proof-of-concept of the possibility of independent detection of cell death of different cell lines in cocultures. Currently, there are not many methods for detecting cell death in 3D cultures, and, as far as the authors know, there are no methods for assessing cell death in cocultures that would allow estimating the percentage of death of each of given lines included in the coculture. Moreover, the described method can be used not only for the purposes of screening promising drugs - the method can be used in modeling pathological conditions when endogenous metabolites produced by one type of cells can negatively affect another type of cells. Quite interesting results obtained as a result of this work include the identified difference in sensitivity to capecitabine when culturing cells in spheroids and in alginate hydrogel. We found differences in sensitivity to cytostatics between 3D cultures in which assembly processes predominate (spheroids) and cultures in which growth and division processes predominate (alginate hydrogel). Currently, this field of in vitro toxicology is virtually unexplored - anecdotal findings include the result of the Senkowski group, who discovered a difference in sensitivity to nitazoxanide when culturing cells in 2D monolayer and 3D spheroids[27].

## Conflict of interest

The authors declare no conflict of interest.

## Funding

This scientific project was financed on the basis of “The Agreement on the provision of subsidies for financial support of the execution of the state task for the provision of public services (performance of work) No. 141-03-2024-014 from January 18, 2024 - Development of molecular genetic diagnostic methods for the quantification of sanogenesis in healthy people (norm) 122030900062-5”

## Notes

### Competing Interest Statement

The authors have declared no competing interest.

